# Differential modulation of heat inducible genes across diverse genotypes and molecular cloning of a sHSP from Pearl millet [*Pennisetum glaucum* (L.) R. Br.]

**DOI:** 10.1101/2020.02.26.966184

**Authors:** S MukeshSankar, C. Tara Satyavathi, Sharmistha Barthakur, S.P Singh, Roshan Kumar, K.V. Prabhu, C. Bharadwaj, S.L. Soumya

**Affiliations:** Division of Genetics, ICAR-Indian Agricultural Research Institute, New Delhi, India; Project Coordinator - Pearl Millet, ICAR-All India Coordinated Research Project on Pearl Millet, Mandor, Jodhpur, India; ICAR-National Institute for Plant Biotechnology, New Delhi, India; National Institute of Plant Genome Research, New Delhi, India; Chairperson, Protection of Plant Varieties and Farmers’ Rights Authority, Government of India, India

**Author notes:** Corresponding Author: (MSS).

## Abstract

Environmental stresses negatively influence survival, biomass and grain yield of most crops. Towards functionally clarifying the role of heat responsive genes in Pearl millet under high temperature stress, the present study were carried out using semi quantitative RT- PCR for transcript expression profiling of *hsf and hsps* in 8 different inbred lines at seedling stage, which was earlier identified as thermo tolerant/susceptible lines through initial screening for thermo tolerance using membrane stability index among 38 elite genotypes. Transcript expression pattern suggested existence of differential response among different genotypes in response to heat stress in the form of accumulation of heat shock responsive gene transcripts. Genotypes WGI 126, TT-1 and MS 841B responded positively towards high temperature stress for transcript accumulation for both *Pgcp 70* and *Pghsf* and also had better growth under heat stress, whereas PPMI 69 showed the least responsiveness to transcript induction supporting the membrane stability index data for scoring thermotolerance, suggesting the efficacy of transcript expression profiling as a molecular based screening technique for identification of thermotolerant genes and genotypes at particular crop growth stages. As to demonstrate this, a full length cDNA of *Pghsp 16.97* was cloned from the thermotolerant cultivar, WGI 126 and characterized for thermotolerance. The results of demonstration set forth the transcript profiling for heat tolerant genes can be a very useful technique for high throughput screening of tolerant genotypes at molecular level from large cultivar collections at seedling stage.

## Introduction

Pearl Millet [*Pennisetum glaucum* (L.)] R.Br. is an annual warm season C_4_ cereal crop, widely cultivated in the semi-arid tropical (SAT) regions of Africa and the Indian subcontinent covering an area of 29 million ha, forms staple food and fodder for 90 million resource poor inhabitants [1]. India is the largest producer of this crop among world grown over an area of 7.38 million ha with a production of 9.13 million tons during 2018-19 [2]. Pearl millet is widely taken up as kharif crop with least input during the peak summer periods with the onset of monsoon. High temperature spells beyond 42°C accompanied by moisture stress during seedling stage of the crop especially during the germination and seedling establishment stages will affect the adequate plant population. It continued to be a severe constraint to Pearl millet production under subsistence farming conditions of the semi-arid regions of India and Sub Saharan Africa which ultimately reflects in crop growth and development thereby its productivity in terms of quantity and quality will deteriorate [3]. This situation demands more and more attention not only for the development of stress tolerant genotypes, but also in identification and characterisation of genes responsive for tolerance.

Plants being sessile have the ability to dramatically alter their gene expressions in response to various stress signals [4]through a series of morphological, physiological and molecular alterations that adversely affect plant growth and productivity. Acquisition of thermotolerance is largely controlled through molecular mechanisms based on the activation and regulation of specific stress-related genes. In response to heat stress, Pearl millet produces an array of proteins which helps in alleviating the effects of stress. One such major protein family are heat shock proteins (HSPs). HSPs/chaperones have been found to play a role in stress signal transduction and gene activation [5]. Heat stress-response signal transduction pathways and defence mechanisms, involving HSFs (Heat Shock Factors) and HSPs, are reported to be involved in the sensing of Reactive Oxygen Species (ROS) [6]. HSP transcripts were shown to be helpful in diagnosing plant stress [7] and considerable evidence for specific heat shock proteins involved in the development of thermotolerance in Pearl millet was first reported by Howarth [8]. Based on their approximate molecular weights, five major families of Hsps are recognized: Hsp100, Hsp90, Hsp70, Hsp60, and the small Hsp (sHsp) families [9]. Among which HSP 90, 70 and sHSP were well studied in Pearl millet [10,11,12], suggested cytosolic Hsp70 and sHSPs are widely associated with thermotolerance in germinating seedling [13] and maintenance of cell membrane fluidity under high temperature stress [14] which were the two major indices [15,16] used for screening seedling thermotolerance practically. HSP 70 functions as molecular chaperones for nascent proteins by proper folding and prevention of their accumulations as aggregates, its transport to their final location and also plays a pivotal role in disassembly of non-native protein aggregates and their subsequent refolding and recovery from stress-induced protein damage [10]. Small heat shock proteins (sHsps) are diverse groups of proteins that are conserved in both eukaryotes and prokaryotes with molecular weights in the range of 15–40 kD, whose expression were limited in the absence of environmental stress, got up-regulated to over 200 fold upon induction of heat stress [17]. It play a critical role in organismal defense during physiological stress as they protect proteins from irreversible aggregation by an energy-independent process until suitable conditions pertain for renewed cell activity, at which time protein release and refolding are mediated by ATP-dependent chaperones such as Hsp70 [18]. A recent report concluded that there were some indications that small heat shock proteins play an important role in membrane quality control and thereby potentially contribute to the maintenance of membrane integrity especially under stress conditions [19].

The transcription of these genes were under control of a master regulatory proteins called heat shock transcription factors (Hsfs) which acts as transcriptional activators for heat shock [20]. It has a major role in coordination of regulatory functions in different stages of response to periodical heat stress such as triggering, maintenance, and recovery. Induction of many heat-inducible genes is attributed to the conserved heat-shock element (HSE) in the promoter. HSE consists of alternating units of pentameric nucleotides (5’-nGAAn-3’) that serve as the binding site for heat shock factor (HSF). In response to heat stress, HSF is converted from a monomeric to trimeric form in the nucleus and was targeted towards concern HSP gene where it has a high affinity of binding to HSEs. It is believed that interaction of HSF with HSP70 or sHSPs results in the activation of transcription of these genes. There are several report which corroborated the overexpression of these protein coding gene enhanced the thermotolerance among plants such as Pearl millet HSP 70 and 90 [10], Rice HSP 70, overexpression of SlHsfA3 in Arabidopsis plants, isolated from cultivated tomato *etc* [21].

Howeverthe picture of molecular mechanism of thermo-tolerance is highly complex consists of poorly understood, various interlinking gene networks [22]. Henceit became a felt need to draw the detailed aspects of the correlated gene expression activities and identifying those genes or gene product which have maximum potency in imparting thermo-tolerance among genotypes at various crop growth stages, for which gene based genotype screening come to be inevitable. The above said facts more significant as the threat of climate change and global warming become a matter of reality as according to the report of the United States Environment Protection Agency, the global average temperature has risen by about 1.4°F in the past century and is expected to increase by 2°F to 11.5°F by 2100 [23]. Rising temperatures may lead to altered geographical distribution and growing season of agricultural crops by allowing the threshold temperature for the start of the season and crop maturity to reach earlier [24]. There is a constant need for the identification, isolation and characterisation of increasing number of stress induced genes unravelling their functions for enhancing agricultural productivity.

A major challenge in conventional breeding for heat tolerance is the identification of reliable and effective screening methods to facilitate detection of heat-tolerant plants and the genes responsible for thermo-tolerance. In present study, we performed semi quantitative RT-PCR (Reverse Transcription Polymerase Chain Reaction) for expression profiling of *Pennisetum glaucum heat shock factor* (*Pghsf)* and *Pennisetum glaucum chloroplast localized HSP 70 (Pgcp70)* transcripts in different cultivar at seedling stage, screened for thermotolerance. The correlated expression pattern of *Pghsf* and *Pgcp70* were also studied which revealed the initiator role of *Pghsf* and early response role of *Pgcp70* towards high temperature stress. We also carried out isolation and characterization of full length transcript of *PgHSP16.97*, a gene encoding alpha-crystalline sHSP shows specific expression patterns during water and high temperature stress. This paper give validation on the effectiveness of RT-PCR based screening methods for the identification and utilisation of thermotolerance genes from superior heat tolerant genotypes for bridging supra-optimal temperature tolerance with high productivity in Pearl millet.

## Material and methods

### Plant material

Eight elite Pearl millet inbred lines (6 thermotolerant and 2 thermosusceptible), earlier screened for thermotolerance (Table 1) using the physiological parameter, Membrane Stability Index [25,26,27] were collected from the Pearl millet breeding unit of Indian Agricultural Research institute, New Delhi (IARI) and selections made from International Crop Research Institute for Semi-arid Tropics, Hyderabad (ICRISAT) and Central Arid Zone Research Institute, Jodhpur (CAZRI) were used for expression studies.

**Table 1.**
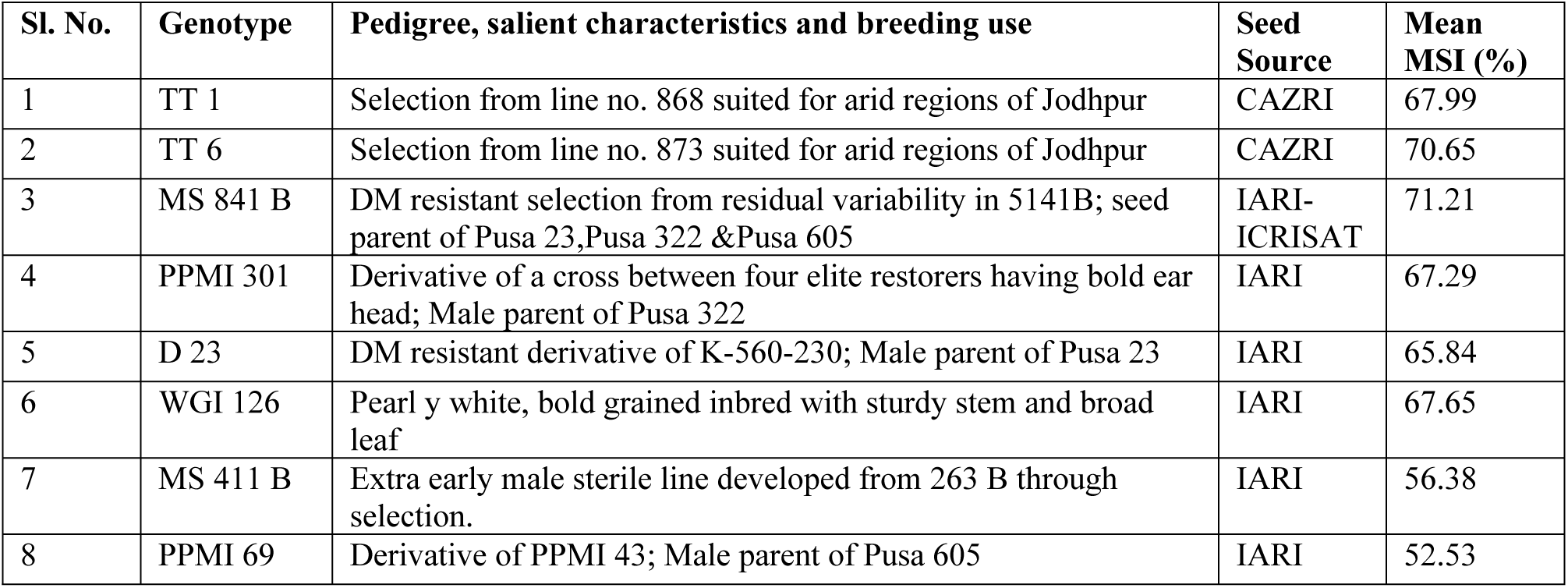
Details of genotypes used for transcript expression studies for heat tolerance in Pearl millet.

### Temperature treatment

Seeds were grown in a pot containing autoclaved soilrite were kept under constant light/dark regime with 16 hrs (hours) light and 8 hrs darkness at 25°C in culture room were used for expression studies. Heat stress was imposed on 7 and 10 day old seedlings in a growth chamber at 42°C for 2hrs and 6hrs respectively before tissue harvest. For comparison, seedlings grown at regular temperature in culture room were used as control (Fig 1).

**Fig 1:**
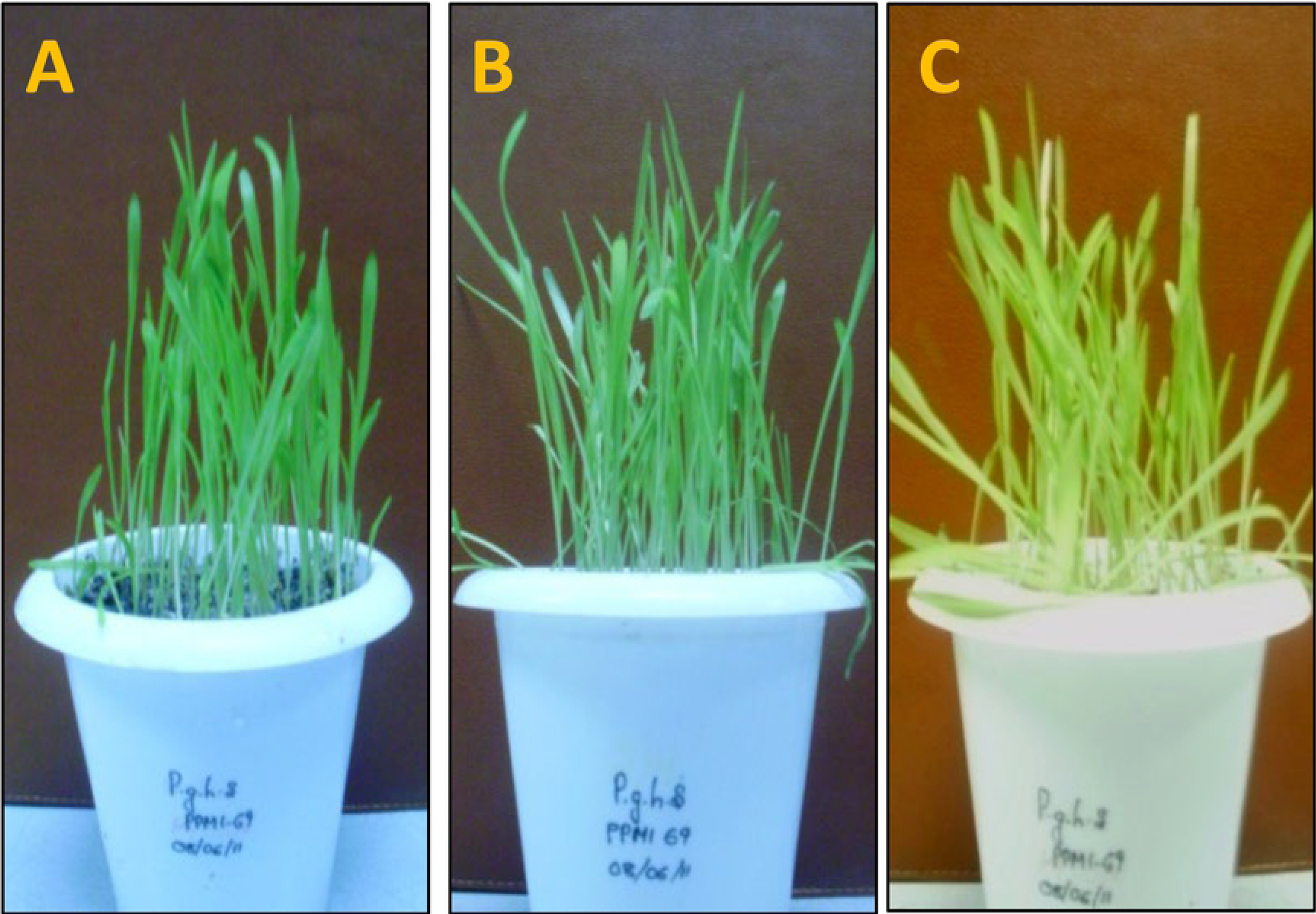
Response of Pearl millet seedling to control (regular growth condition) and severe stages of the heat stress (42°C) used for expression analysis (A: Control 25°C, B: 42°C for 2Hrs & C: 42°C for 6Hrs)

### Sample collection

Aerial portion of seedlings were harvested after heat treatment given in growth chamber. The samples were also collected from plants raised in culture room, kept under controlled conditions. Leaf samples were harvested using sterile scissors, wrapped in aluminum foil, labeled and then immediately transferred to liquid nitrogen. The samples were then taken for storage in −80°C freezer kept in lab, National Institute for Plant Biotechnology, New Delhi.

### Isolation and quantification of total RNA

Total RNA was isolated by Triazol method (Invitrogen, Carlsbad, CA, USA) followed by quantification using Nano Drop spectrophotometer (ND 1000, Thermo Scientific, USA) as well as by Agarose Gel Electrophoresis. DNase treatment (Invitrogen, USA) was carried out to remove any contaminating DNA followed by RT-PCR.

### RT-PCR expression analysis of target genes

Coding sequences for two candidate genes of Pearl millet *Pgcp70* (acc. no. EF495353.1) and *Pghsf* (acc. no. EU492460.1) were downloaded from NCBI (http://www.ncbi.nlm.nih.gov) public database. For Semi quantitative RT-PCR expression analysis, primers were designed to specifically amplify the selected mRNA sequence of the above genes maintaining stringency and specificity. The details of the primers with its melting temperatures (Tm) are shown in Table 2.

**Table 2.**
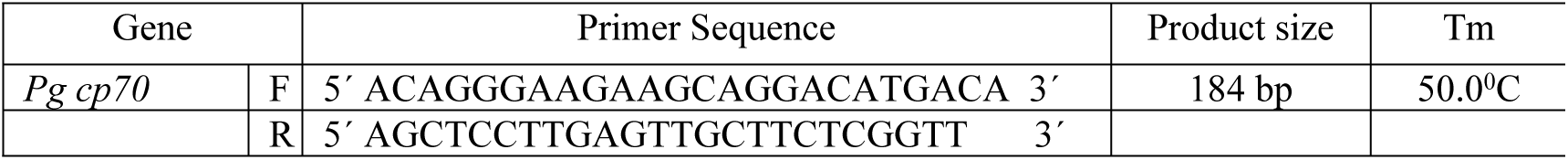
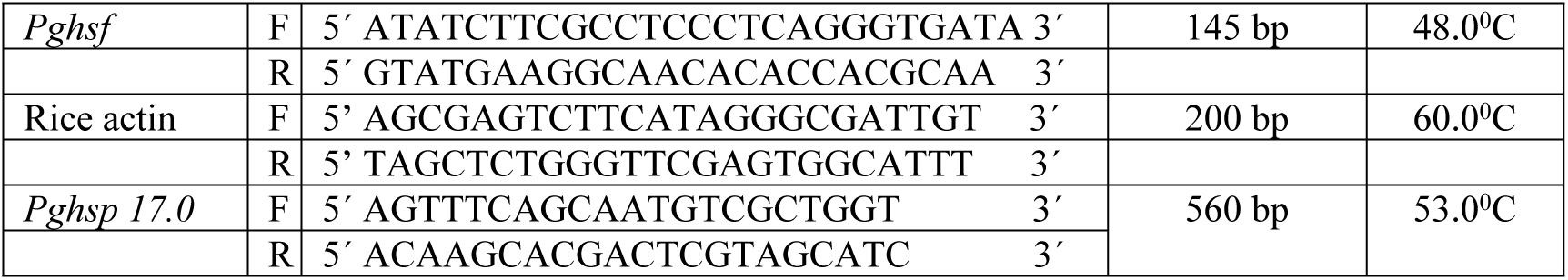
Details of primers used for transcript expression profiling and full length cloning.

Reverse transcription-polymerase chain reaction (RT-PCR) was carried out in two steps as per the protocol. Synthesis of cDNA was conducted using the Thermo, Verso™cDNA-kit, USA (Thermo Fisher Scientific Inc, USA) according to the supplier’s instructions using oligo (dT)_15_ as primer. For expression analysis, cDNA pool (2µL) was used as a template, to amplify the corresponding heat responsive gene transcript by PCR using PCR master mix (PromegaCorporation, USA) as per manufactures instruction. Thirty cycles of PCR (with 4 min of initial denaturation at 94°C, 94°C for 45 sec, 48-60°C (Tm optimized for the individual genes) for 45 sec, 72°C for 1 min) amplification, with a final extension at 72°C for 10 min was performed.

The RT-PCR products were loaded on a 1.2% agarose gel and the stained DNA products were photographed using Alfa Imager gel documentation system HP (Proteinsimple, USA). House-keeping gene, Actin, was used in all expression studies and treated as reference gene (internal constitutive control) to show equal loading and to ensure the integrity of c-DNA, which showed equal expression in all genotypes under various degree of heat stress. The transcript level of each test targets were averaged for triplicate reactions. The gene expression data were normalized by subtracting the mean expression level from reference gene. The relative fold change in expression in treatments (T) was compared with those from regular growth stage (C) was done by expression value of control as calibrator for respective genotype using Alfa Imager Software tools by keeping the density of bands in control as unity.

### Isolation and cloning of a full length Pg HSP

Based on transcript expression profiling studies, the best thermotolerant genotype was used to isolate a full length cDNA of one small heat shock protein *Pghsp17.0* (Acc. No. X94191.1). Primers were designed (Table.2) to carried out RT-PCR as described above and the fragment obtained was purified and sequenced. After sequencing and confirming the isolated amplicon as *Pghsp17* and the purified cDNA was cloned onto pGEM-T vector (Promega, USA) through TA cloning and transformed into *E.coli*-XL1 blue competent cells. Based on blue white screening, ampicillin resistance putative recombinants were selected for further analysis by colony PCR. Positive clones were inoculated overnight in LB (Luria–Bertani Agar) and the plasmid was isolated. Restriction digestion of plasmid DNA with ECoRI in Takara RE kit (Clontech Bio Inc, Japan) was done to further confirm successful cloning of *Pghsp17.0 gene* from Pearl millet cultivar WGI 126.

### DNA sequencing and data analysis

Full length c-DNA fragment were isolated and sequenced at the Xcelris Labs Ltd, Ahmedabad, India. Analysis of the c-DNA sequences was performed using the BLASTn program [28]. The conceptual translation of nucleotide sequence was made using the Expasy translate tool. (http://web.expasy.org/translate/). Multiple sequence alignments were carried out using the CLUSTALW software package [29] and thus phylogenetic analyses were performed with all full-length *HSP 17.0* protein sequences publicly available for *Pennisetumglaucum, Zea mays, Oryza sativa, Triticumaesativum, Sorghum bicolor* and *Hordeumspp*usingthe CLUSTALW program in MEGA 5.2 software and created phylogenic tree by neighbour joining method after bootstrapping for 500 times using previously aligned amino acid sequences.

### Homology modeling and structure analysis

Three dimensional structure of *Pg HSP 17.0* was deduced by Modeller v9.11 [30], was subjected to backbone conformation evaluation by investigating psi/phi Ramachandran plot using Procheck[31]. The final model and the template were subjected to superimposition for structural comparison using STRAP interface (http://www.bioinformatics.org/strap/).

## Results

### Relative transcript expression profiling of Hsp and Hsf under high temperature stress

The cDNA were synthesized from mRNA isolated from heat stress (42°C) exposed *Pennisetumglaucum* seedlings of six tolerant and two sensitive genotypes. Further these c-DNA were used for expressions profiling studies of two high temperature responsive genes namely *Pgcp70* and *Pghsf* by semi quantitative end point RT-PCR. The expression pattern were analyzed based on visual analysis of gel and their densitometric semi quantitative quantification using alpha imager software and were used to understand their correlated expression pattern under high temperature stress.

Transcript expression profiling for *Pgcp70* showed differential expression pattern under regular and high temperature stresses during the time course of experiment among different Pearl millet genotypes (Fig 2A). Even though *Pgcp70* got expressed under regular growth, the expression level of *Pgcp70* got increased very significantly upon heat stress. The genotypes showed a significant variability for transcript accumulation upon heat stressin which, the level was up-regulated in thermotolerant lines, while in susceptible genotype (PPMI 69), the gene got down regulated by 38% and in MS 411B the expression was comparatively less. The expression profiling of *Pgcp70* suggested the HSP 70 was highly induced at early stage of heat exposure (for 2 hrs) whose transcript level was slightly increased as heat stress progressed for long duration of 6 hrs on 7 day old seedlings. Even though we observed constant induction of HSP 70 during heat stress on 10^th^ day old seedling, the result suggested that heat stress during early phase (for 2 hrs) leads to up-regulation of *Pgcp70* transcript in all genotypes while its level diminished with continuous exposure to heat stress (6 hrs) over a period. Also transcript accumulation shows slight increase from 7^th^ day to 10^th^ day old seedling, shows plant tend to increase tolerance to high temperature with growth and development and continued exposure. Among thermotolerant genotypes WGI 126, TT 1, TT 6 *etc* were shown to respond positively to heat stress by showing a relatively elevated level of high temperature induction of *Pgcp70* RNA levels whereas genotype PPMI 69 and MS 411B showed the least induction. The results (Fig 2B) revealed that WGI 126 showed elevated expression of *Pgcp70* 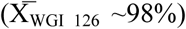 under high temperature particularly 2hrs of heat stress at 10DAS as compared to control plants. The genotype wise high temperature tolerance expression pattern of stress inducible gene *Pgcp70* can be categorised as WGI 126>TT 1>TT 6>MS 841 B>PPMI 301>D 23>MS 411 B>PPMI 69.

**Fig2.**
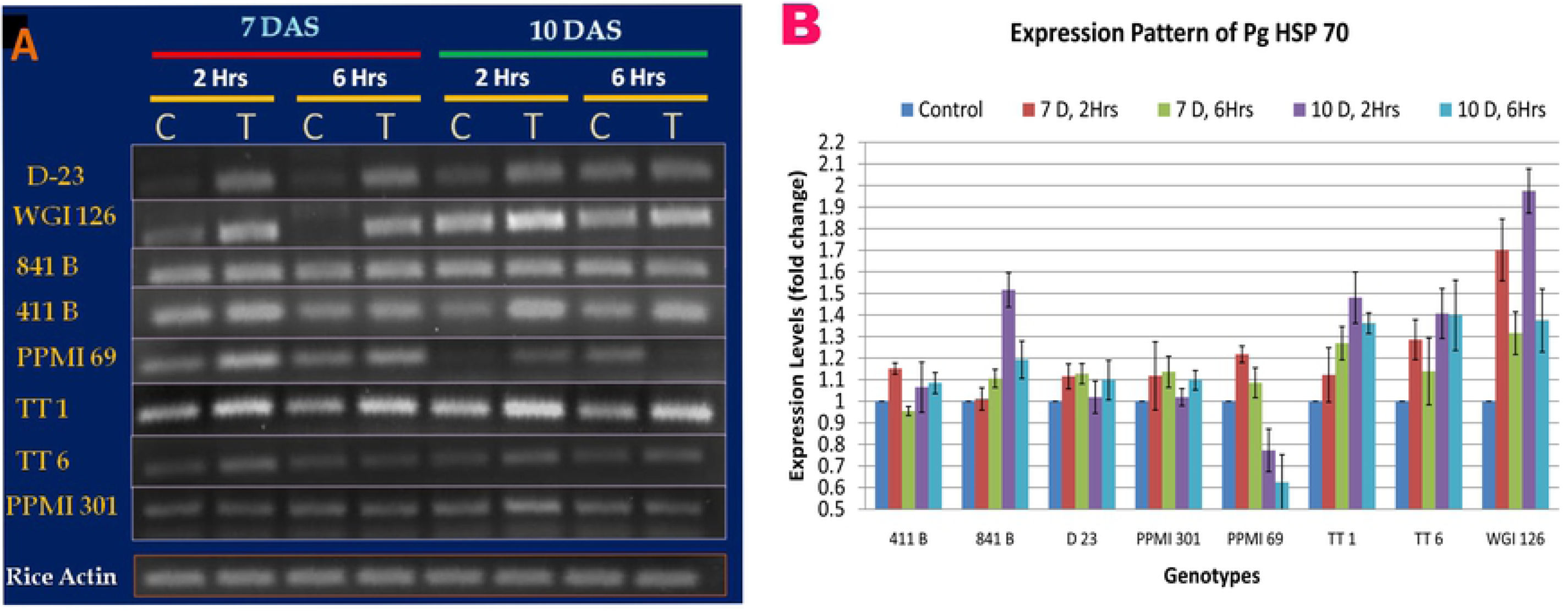
Semi quantitative real time expression profiling of heat responsive gene *Pgcp70* under heat stress of 42°C in 8 selected genotypes of Pearl millet at different growth stages against differential heat treatment. (Fig 2A: densiometric expression pattern among genotypes, observed in 1.2% agarose gel along with Os-actin gene expression was used in Pearl millet as endogenous control to normalise the expression. Fig 2B: Fold change expression level of transcript among genotypes at different stress conditions)

Similar to the expression pattern for *Pgcp70, Pghsf* mRNA also showed differential expression pattern among different Pearl millet genotypes (Fig 3A). Genotypes WGI 126, TT 1 and MS 841B showed elevated level of transcript accumulation on exposure to heat stress and genotype PPMI 69 showed least accumulation of transcript on exposure to heat stress which supported the MSI (Membrane Stability Index) studies conducted earlier [25]. A comparative expression profiling study among thermotolerant genotype to find best thermotolerant genotype using Alfa imager software (Fig 3B) showed higher level of transcript expression (37%) for the genotype WGI 126, 2 hrs of heat stress on 10^th^ day old seedling. Genotypes can be categorised as per *Pghsf* transcript abundance during the expression profiling as WGI 126>TT 1>MS 841 B>PPMI 301>TT 6>D 23>MS 411 B>PPMI 69.

**Fig 3.**
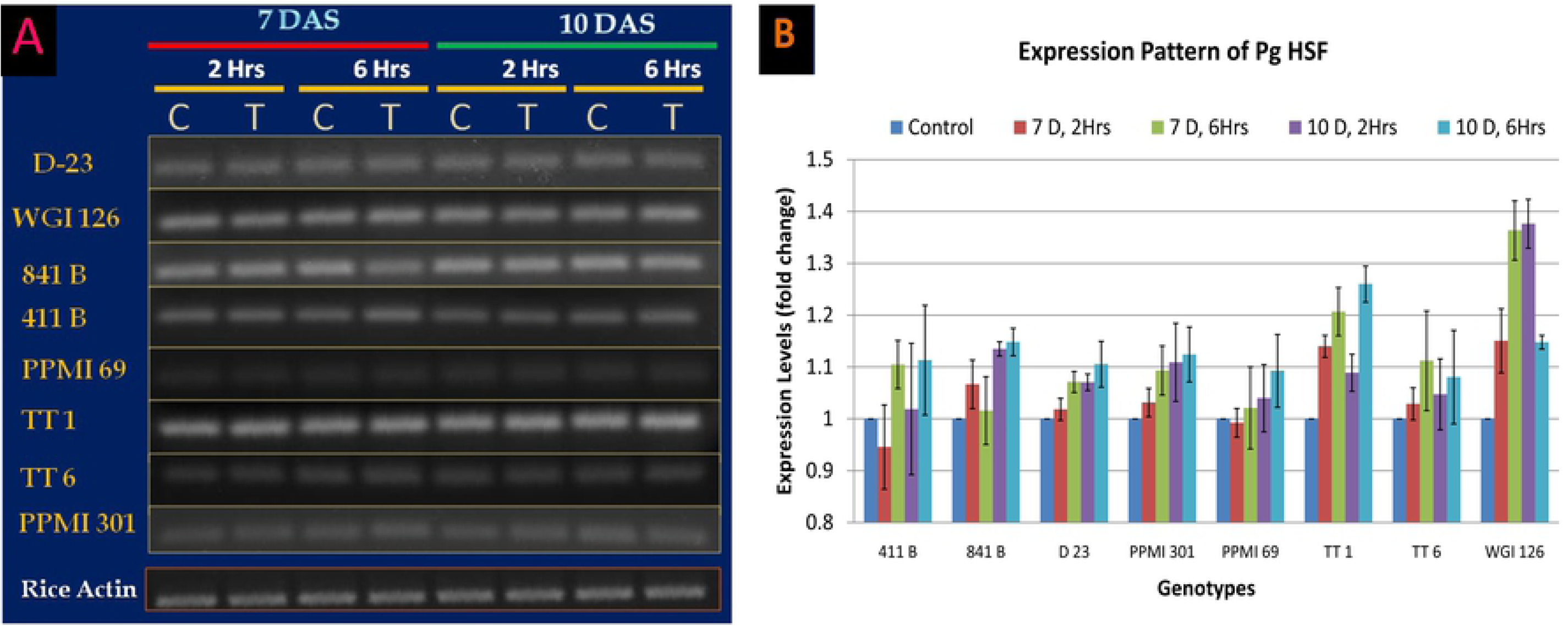
Semi quantitative real time expression profiling of heat responsive gene *PgHSF* under heat stress of 42°C in 8 selected genotypes of Pearl millet at different growth stages against differential heat treatment (Fig 2A: densiometric expression pattern among genotypes, observed in 1.2% agarose gel along with Os-actin gene expression was used in Pearl millet as endogenous control to normalise the expression. Fig 2B: Fold change expression level of transcript among genotypes at different stress conditions)

Comparative expression studies between two genes *Pghsf* and *Pgcp70* among genotypes (Fig 4) suggested, even though *Pghsf* showed lower expression under induction of heat stress during initial phase (2hrs), its expression got a steady increase (4.6% to 12.3% and 11.0% to 13.3% at 7 & 10D old seedling respectively) upon prolonging the heat stress upto 6hrs. In contrast to *Pghsf, Pgcp70* had relatively faster response kinetics and reached its peak expression at early stages of heat stress (2hrs) and its level diminished (21.5% to 14.2% and 28.2% to 15.4% at 7 and 10D old seedling respectively) after a prolonged treatment of 6hrs. The transcript expression upon heat stress increased for corresponding differential heat treatment as the age of seedling progressed, as the 10D old seedling shown to have more accumulation of transcript than that in 7D old seedling.

**Fig 4:**
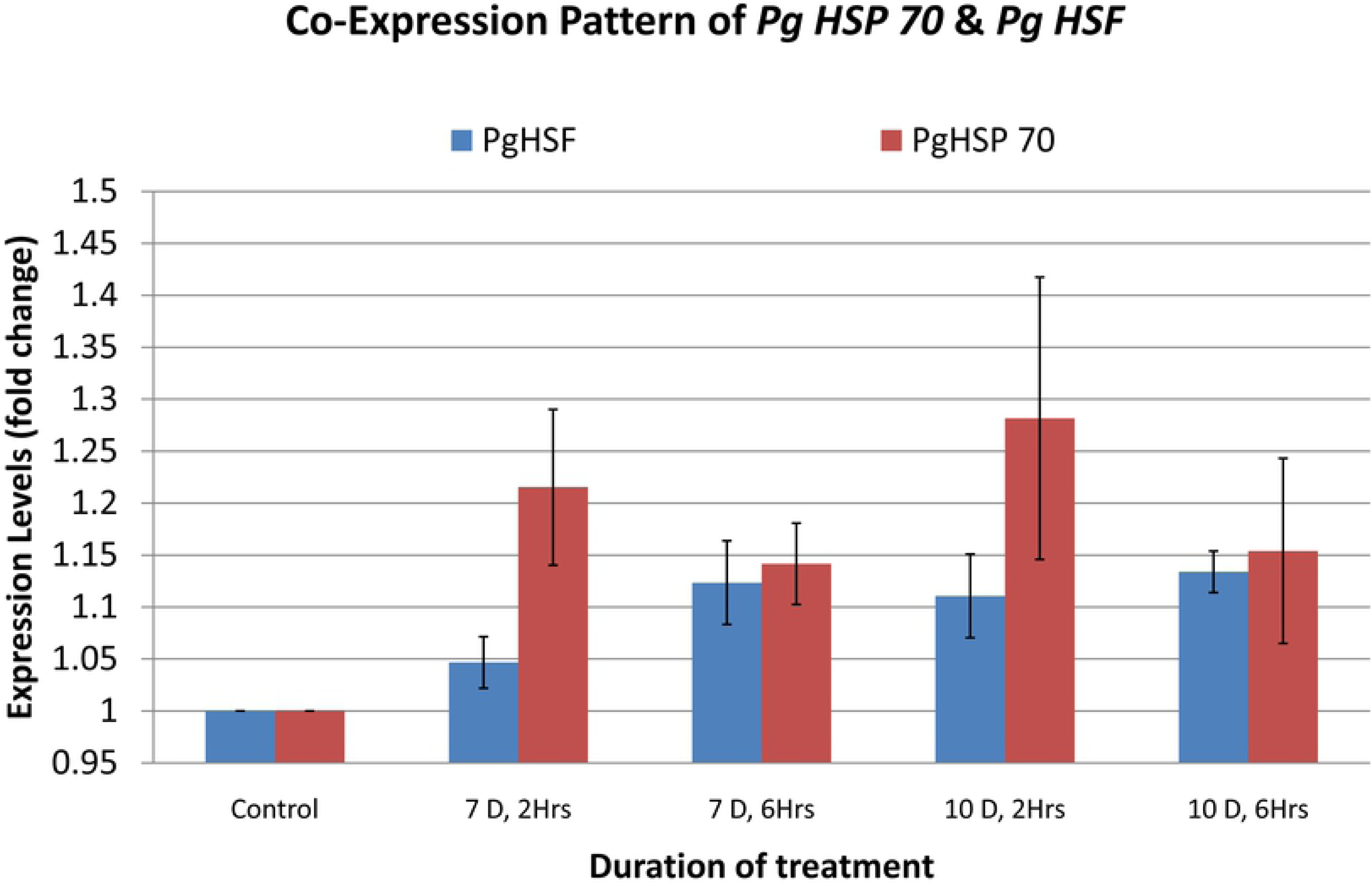
Co-Expression Pattern of *Pgcp 70* &*Pg HSF* of Pearl millet under differential heat stress.

### Isolation and Cloning of full length cDNA of *Pg HSP 17.0* from Pearl millet

A c-DNA of small heat shock protein family member of approximately 600bp was isolated from the tolerant cultivar WGI 126 after exposing to high temperature stress at 42°C for 6 hrs by RT-PCR using *Pg HSP 17.0* specific primers. The fragment was excised from gel, purified and cloned in TA cloning vector, pGeMT easy (Promega) and positive clones were selected by blue white screening (Fig 5). They were further confirmed by colony PCR using *Pg HSP 17.0* specific primers. Positive colonies were inoculated in LB supplemented with ampicillin overnight and plasmid DNA isolated. Restriction digestion was carried out with the enzyme EcoRI which release the insert from the vector (Fig 6).

**Fig 5.**
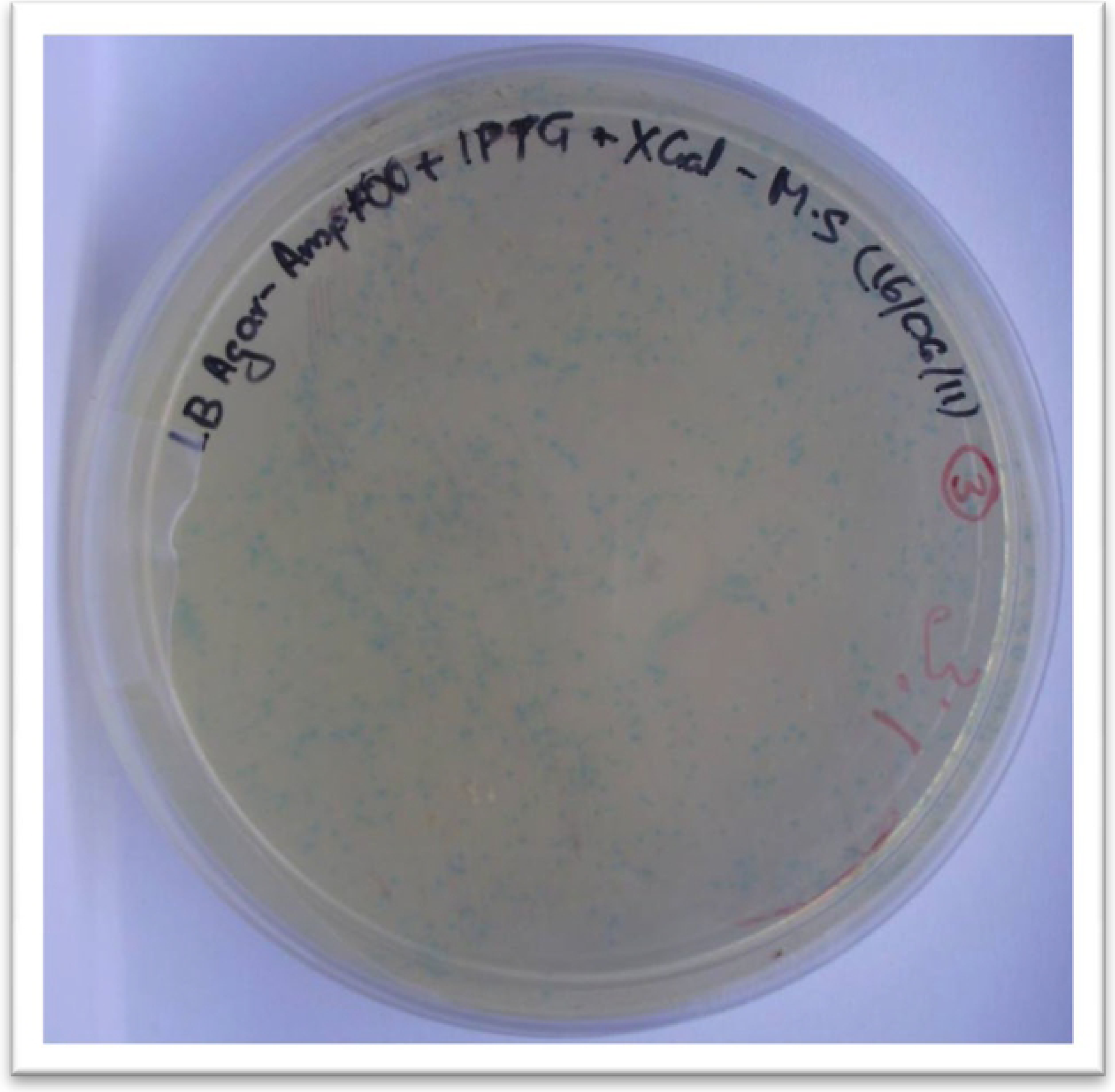
Selection of Recombinant pGM-T*-PgHSP17 E. coli cells* by Blue white screening of colonies.

**Fig 6.**
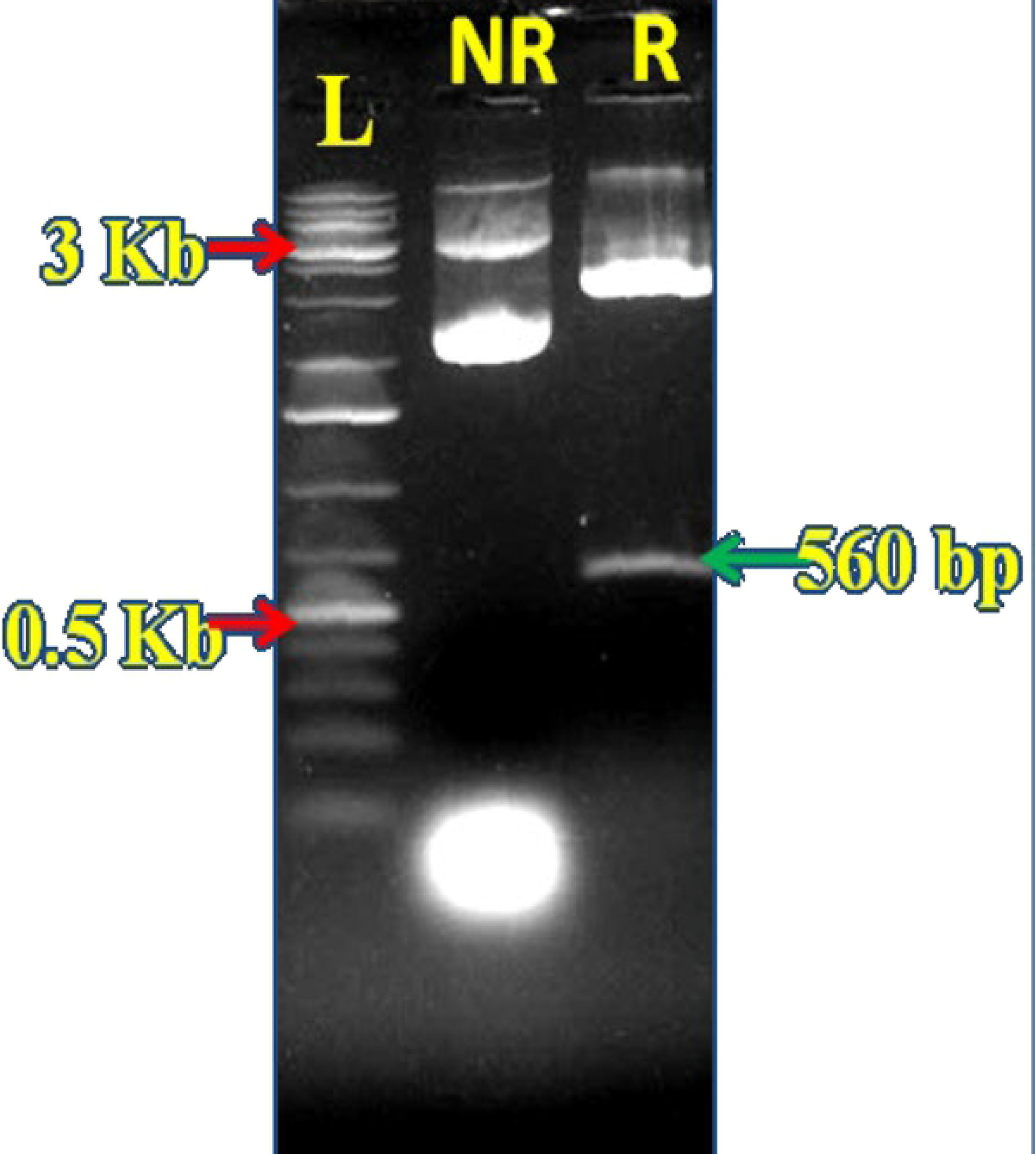
Restriction analysis of non-recombinant (NR) and recombinant (R) cloned by EcoR I shows the upper band (3 Kb) corresponds to vector DNA and lower band (560bp) corresponds to *PgHSP17* insert DNA

The cDNA inserts of these recombinant plasmids were sequenced completely. The sequence analysis of RT-PCR product (Fig 7a) using BLAST program suggested theisolated fragment had one full length cDNA with single ORF (Open reading frame) with a size of 560bp having close similarity with *Cenchrushsp 17.0*. (Acc. No:X94191). The full-length cDNA of PgHsp17.0 have a size of 560bp which contained an open reading frame of 459 and 28bp 5’ and 73bp 3’ untranslated regions (UTRs). The translation initiation region of this open reading frame was situated within a sequence, CCATGG, which resembled the plant consensus initiation sequence [32], but the consensus polyadenylation signal (AATAAA) was not found in the 3’UTR. In plants, however, repeated AT-rich sequences are regarded as an alternative polyadenylation signals in nuclear genes of higher plants [33].

**Fig 7.**
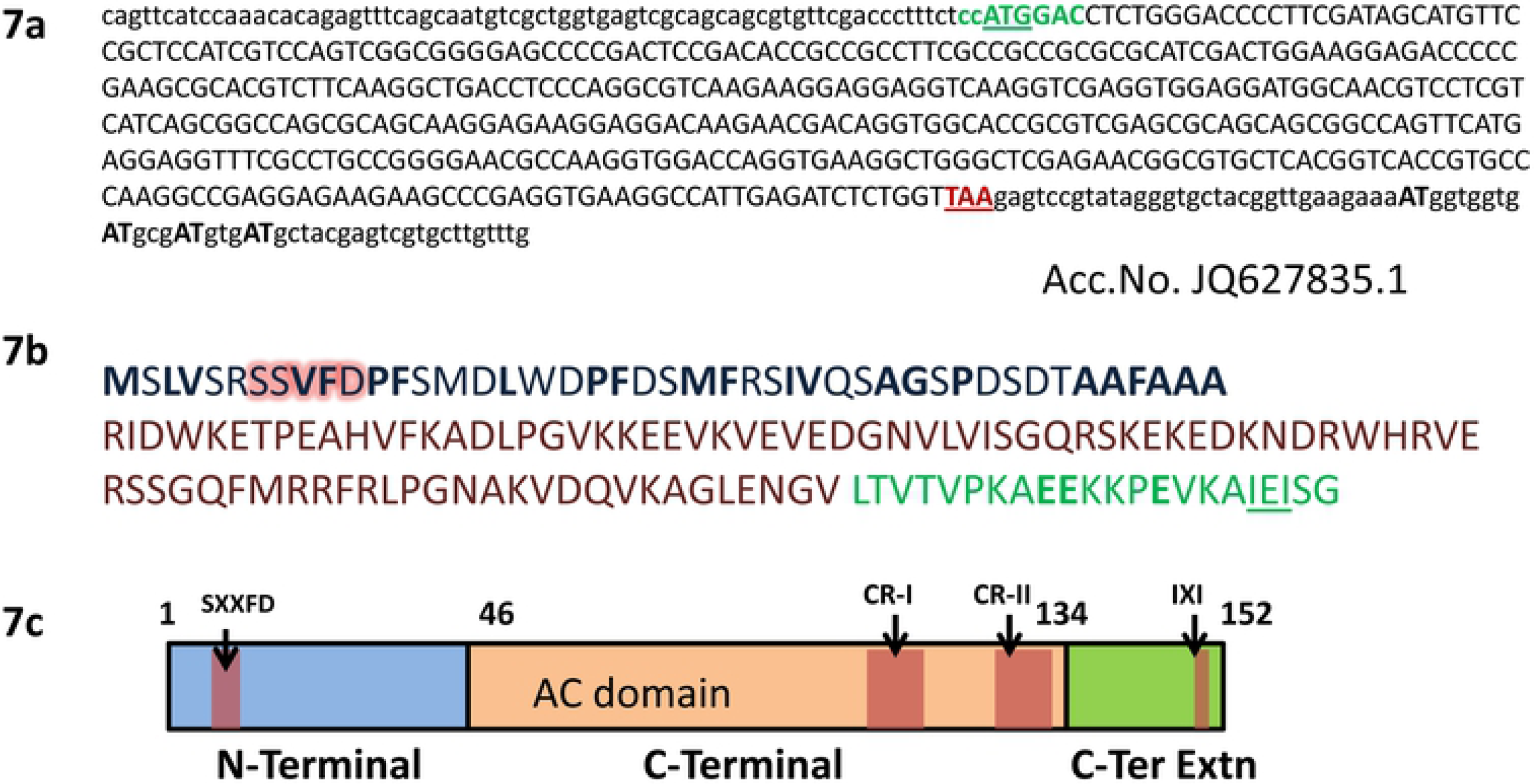
Structural organization of the *PgHsp17.0* gene. **a.** The c-DNA nucleotide sequences, wherein the coding region (upper case letters), 5’, 3’ UTR regions (lower case letters). Translation start site and termination codon are underlined, Polyadenylation signals (repeated “AT” rich sequence given in bold). **b.** Deduced amino acid sequence is placed beneath the c-DNA nucleotide sequences. Various functional domains in the sequence have been significantly marked, such as Variable N-terminal domain (Blue font) with hydrophobic groups represented by Bold letters, alpha crystalline domain (Brown font) and c terminal extension (green font) in which glutamic acid residues (Bold green) and ‘IXI’ residues are represented. **c.** Schematic representation of *PgHsp17.0* protein structure, including three motifs: the N-Terminal (Blue box), Alpha crystalline domain (orange box), and the c-terminal extension (green box).

### Structure of *Pg HSP 16.97* studied by *in silico* analysis

The *PgHsp17.0* ORF encoded for a protein of 152 amino acids with an apparent molecular weight of 16.97 kDa and an estimated isoelectric point of 5.79. It has been named as *PgHSP16.97* following convention. The homology search done using deduced amino acid sequence of *PgHSP16.97* against the translated non-redundant nucleotide database clearly suggested, the *PgHSP16.97* was related to the other eukaryotic sHsps and showed an overall 100–88% sequence identity with sHSPs of Cenchrusamericanus (Acc # CAA63901.1), *Zea mays* (Acc # NP_001150783.1) *Setariaitalica* (Acc # XP_004968025.1), *Saccharum* hybrid cultivar ROC22 (Acc # AFK73383.1). The presence of alpha-crystallin domain (ACD) found in alpha-crystallin-type small heat shock proteins, and a similar domain found in p23 (a cochaperone for Hsp90) and in other p23-like proteins confirmed that the isolated sequence belong to small heat shock protein gene family ProtComp (http://linux1.softberry.com/cgi-bin/programs/proloc/protcomppl.pl) analysis produced integral sub-cellular localization prediction score of 9.9 for cytoplasmic location which indicated *Pg Hsp16.97* belongs to class I sHSP. It also carried a nuclear localization sequence. The Pg HSP 16.97 (Fig 7b) monomer contains a variable N-terminal domain (aa, 1-46), the conserved HSP20 or α-crystallin domain (aa 47-134), and a less variable C-terminal extension (aa, 135-152). The organellar localised sHSPs have the necessary transit, targeting, or signal located on N-terminal of protein which were absent in sequence indicating cytoplasmic localisation. The schematic representation of protein structure with domains are given (Fig 7c).

### Phylogenetic analysis

A phylogenetic study was conducted using 7 cytosolic class I *HSP 17.0* full length protein sequences from different cereals, along with *Pg HSP 16.97* by Clustal W for multiple sequence alignment (Fig 8) followed by construction of phylogenetic trees using NJ method after 500 times bootstrapping using the MEGA 5.2 software (Fig.9). There is considerable diversity in *HSP 17.0* evolution in cereals, with a few conserved motifs and regions such as IXI motifs, Consensus region I (P-X_14-_GVL), Consensus region II (P-X_15-_V-L), R residue at position 114 at C-terminal and SXXFD motif at N-terminal which were the signature regions of cytosolic sHSP. There are only a few highly conserved domains at N-terminus (21/46) observed during sequence alignment among cereal class I sHSP. The *Pg HSP 16.97* is showing close similarity to Pg HSP 17.0 (X94191.1) and is evolutionarily very close to *ZeaHSP 16.9* protein and is more divergent to that of sorghum.

**Fig8.**
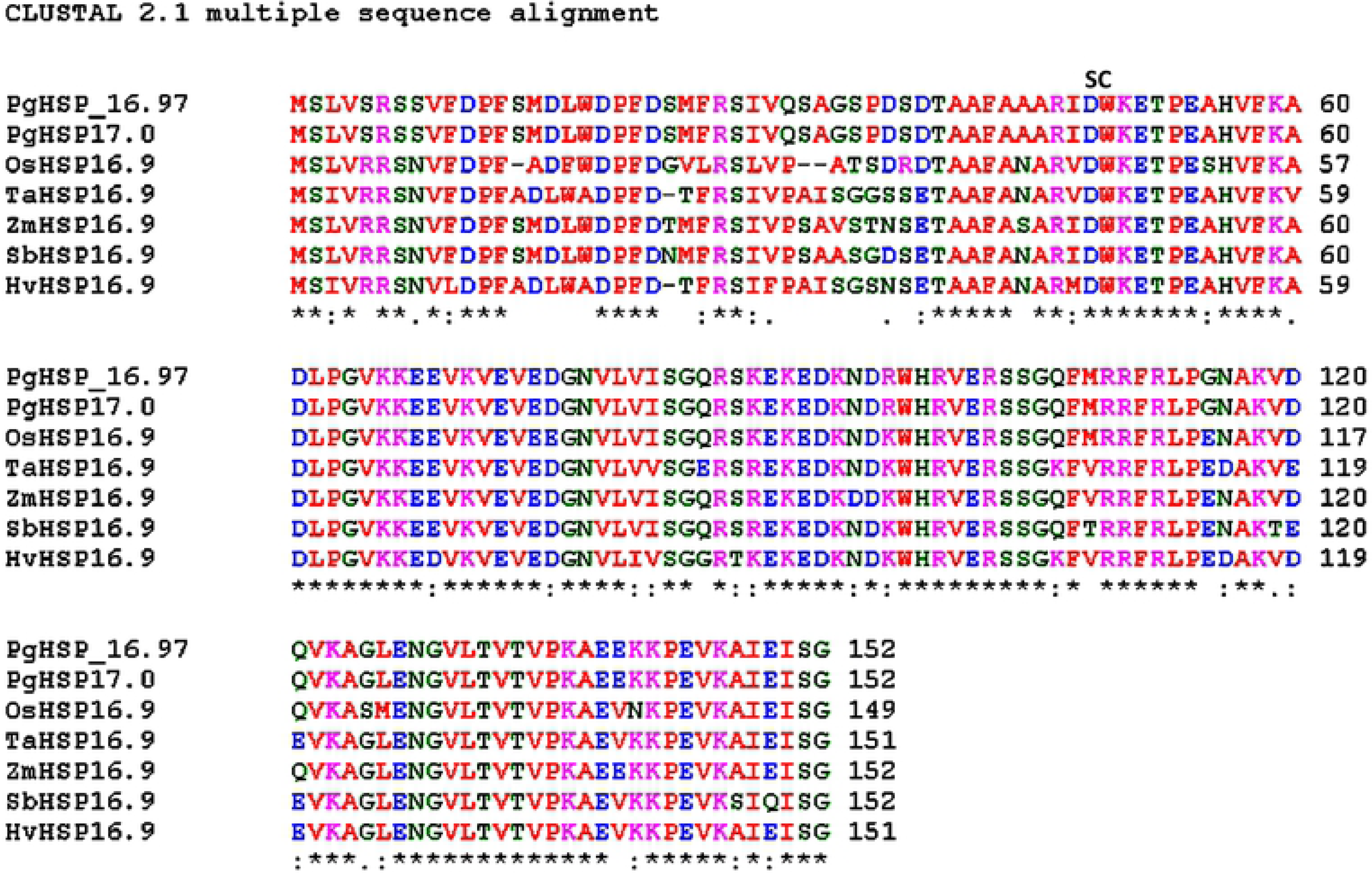
Alignment of cereal HSP 17 Amino Acid Sequences. *PgHsp16.97* (this study), Accession numbers *PgHsp17* (X94191.1), OsHsp (AAB39856.1), TaHSP16.9 (P12810.1), ZmHsp16.9 (ACG24656.1), SbHSP16.9 (XP_002457411.1), HvHSP16.9 (ADW78607.1); SC= start of carboxyl-terminal domain in each alignment, which is the most conserved region of the alpha-crystallin/small heat shock protein (HSP) family; *=conserved residue, := conserved residue with strongly similar property),. = conserved residue of weakly similar property (ClustalW; www.ebi.ac.uk)

### Homology modeling of *PgHSP*16.97

The PgHSP 16.97 and wheat HSP16.9 proteins share 80% similarity at their primary amino acid sequence levels. The crystal structure of wheat HSP16.9 protein (PDB No: 1GME) was chosen as a template for *PgHSP 16.97* model building using the program Modeller 9v11 [30]. Five models of 3D structures of *PgHsp16.97* were generated at various refinement levels were generated and validated using the program Procheck. The best model with a Procheck score of −0.09 was selected. The Accelrys Discovery Studio 3.5 Client program was used to depict the PgHsp16.97 molecular model (Fig 10a). Superimposition of the model with the template and root mean square deviation (RMSD) calculation was done using the program STRAP (http://www.bioinformatics.org/strap/) The RMSD value of the selected *Pghsp 16.97* model structure is 1.60A° with respect to the template 1GME. The structural superimposition was done using STRAP interface observed to have better level of model superimposition onto template which is shown in (Fig 10b).

**Fig 9.**
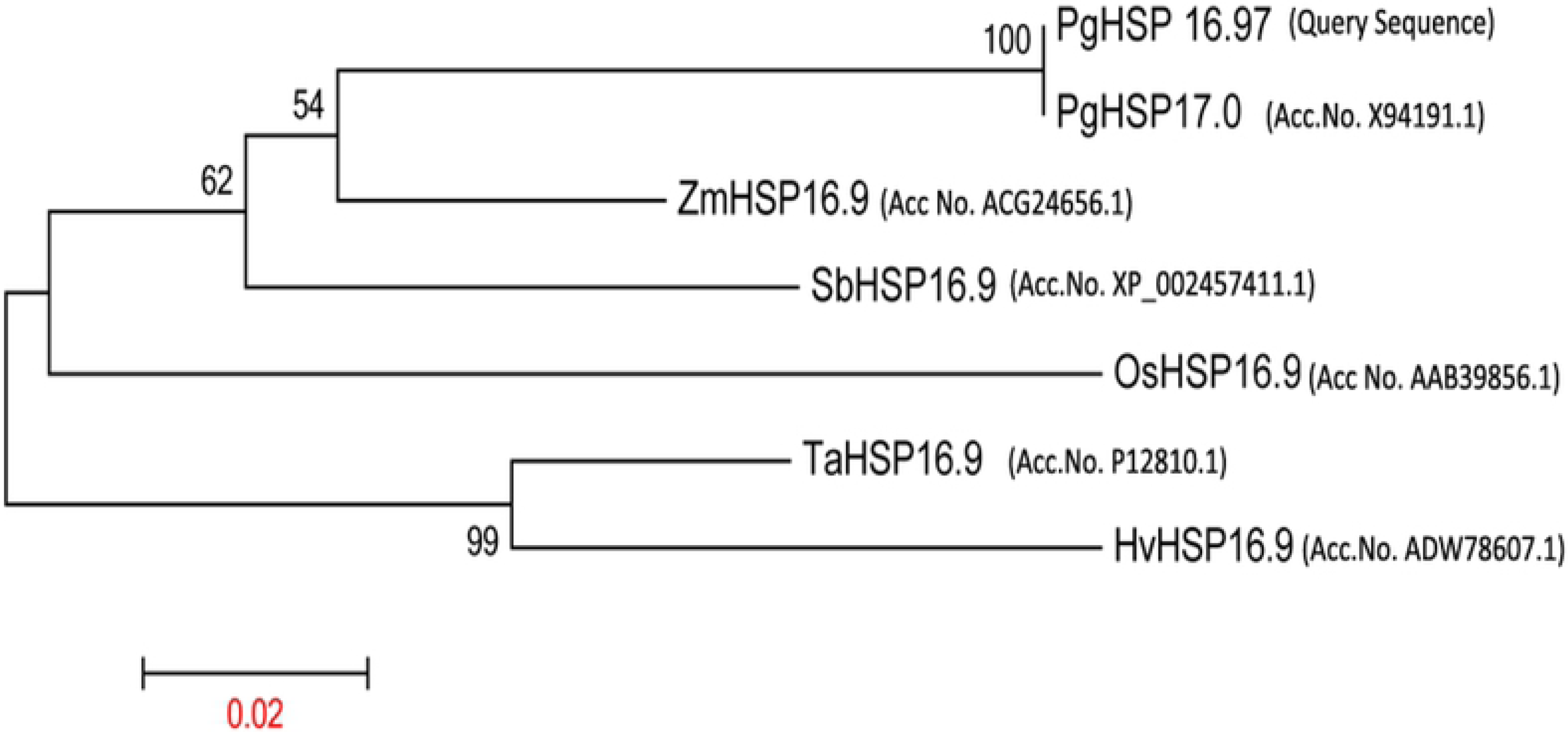
Phylogenetic tree of Cereal *HSP17 by MEGA 5.2*.

**Fig: 10.**
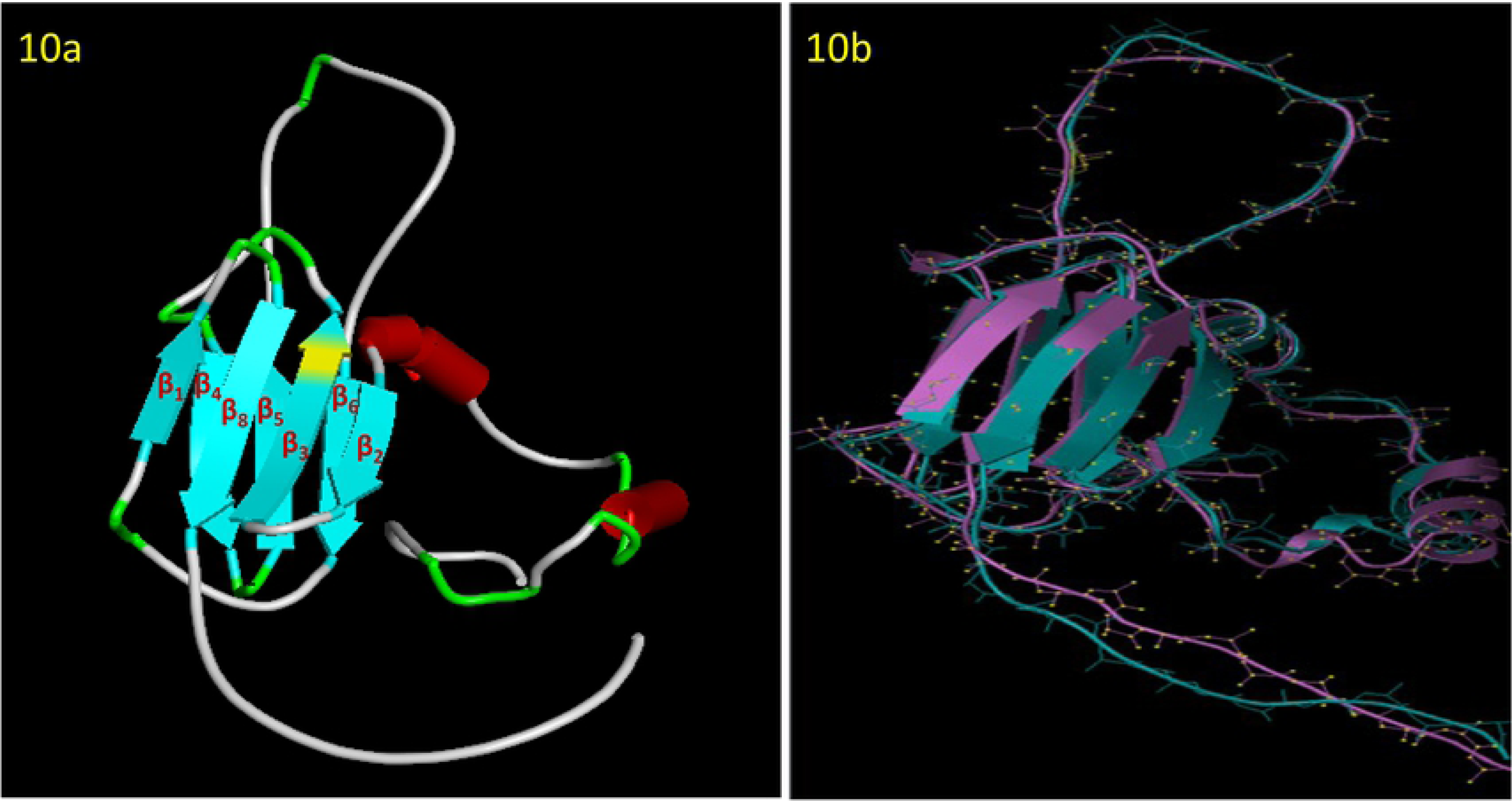
Predicted 3D molecular model of Pg*HSP16.97*. (A: Predicted 3D structure of PgHSP16.97 by modeller 9v11 & B: Structural superimposition between model and template protein (1GME) given by STRAP (Blue: Template and Violet: model)

## Discussion

### RT-PCR expression analysis of HSP and HSF genes

In this study, heat stress lead to induction of various thermotolerant genes in Pearl millet genotypes like Hsfs and HSPs genes, which were evident from the up regulation of transcript level of heat responsive genes such as *Pgcp70* and *Pghsf* under high temperature stress supporting the earlier finding about the heat induced HSPs [34,8,35]. It was also noticed that under normal growth condition (25°C), there was slight accumulation of these transcript, as HSP 70 and its master regulator, hsf have critical role in seed germination and development [36] and also could be attributed to the inherent thermotolerance in Pearl millet cultivars as shown by membrane stability index results. This was correlating with another observation made in Pigeon pea which is widely grown pulse crop in semi-arid regions [37]. The HSP 70 is major molecular chaperon found in all eukaryotes and the gene *PgHsc70* in Pearl milletwas characterised by Reddy *et al*., [10] and it was found to be the heat-shock inducible Hsp70 which is expressed at very low levels under normal conditions, but can be induced rapidly by heat shock and other environmental stresses. The expression profile of *Pgcp70* also generated the same result that *Pgcp70* produced in small quantity at normal condition got enhanced with heat treatment. Large quantity of HSP 70 at early stage of heat exposure (2 hrs), indicates a major role in heat stress during early phases of heat stress. Later on transcript level diminishes as stress is continued (6 hrs) which indicates an immediate shock in genotype leads to higher and rapid induction of *Pgcp70* whose level diminishes with continuous exposure to heat stress suggesting constant involvement of HSP in heat shock for a longer duration as observed in previous studies [38,39]. It was also noticed that *Pgcp70* transcript accumulation shows slight increase from 7^th^ day to 10^th^ day old seedling, shows that plants tend to increase tolerance to temperature as plant develops.

When compared to the transcript level of *Pgcp70, Pghsf* were expressed at low levels. The low level of the hsf transcript was enough to trigger the transcription of *Pgcp70* under heat stress. The transcript expression of *HSF* gene in Pearl millet indicated, its stress regulation initiator role, which showed maximum increase in transcript level in response to heat stress within 30 min of exposure and gradually come down as time proceeded and this result was well matching with the previous study [40]. The expression profile of *Pghsf*studiedwas nearly constant for long period of heat exposure for 2 hrs and 6 hrs and compared to expression profile of *Pgcp70, Pghsf* transcript abundance was less under high temperature treatment in seedling stage than *Pgcp70* suggesting a role in the initial stage rapid stress response in germinating seedling, all further supported the above fact.

There was differential expression pattern for these two heat responsive genes among different genotypes, in which WGI 126 have shown the positive response to heat stress by accumulating more transcript, while PPMI 69 with low accumulation of transcript as expected from the MSI data [26] which suggest the transcript expression profiling can be used for screening thermotolerant and susceptible lines in a large pool of genotypes along with identifying the potential genes responsible for thermotolerance at various stages of crop growth.

### Isolation and Cloning of full length cDNA of *Pg HSP* from Pearl millet

Of the molecular chaperones, the sHSPs were diverse and found in both prokaryotes and eukaryotes, usually undetectable in plant cells under normal physiological conditions, but were induced upon stress lead to plant tolerance to stress, such as drought, salinity, reactive oxygen species, and low temperatures [41]. It is believed that diversification and abundance of the sHsps in a plant reflect an adaptation of the plant to heat stress [9]. Among the sHSPs, the HSP 20 type forms first line of defence against stress in the cell during heat stress will bind to partially folded or denatured proteins, which prevents irreversible unfolding or incorrect protein aggregation, or binds to unfolded proteins by an energy-independent process until suitable conditions pertain for renewed cell activity and allows further refolding by Hsp70/Hsp100 complexes hence been referred to as ‘paramedics of the cell’ [42]. A review concluded that there were some indications that small heat shock proteins play an important role in membrane quality control and thereby potentially contribute to the maintenance of membrane integrity especially under stress conditions [14]. Hence we have isolated and cloned full length cDNA encoding for *HSP 17.0* from *P. glaucum*to understand the structural signature present on this protein for its heat tolerance role. Nucleotide and deduced amino acid sequence analysis of the cDNA clone revealed the presence of alpha-crystallin domain (ACD) found in alpha-crystallin-type small heat shock proteins, and a similar domain found in p23 (a cochaperone for Hsp90) and in other p23-like proteins confirmed that the isolated sequence belong to small heat shock protein gene family. Alpha-crystallin occurs as large aggregates, comprising two types of related subunits (A and B) that are highly similar to the small (15-30kDa) heat shock proteins (HSPs), particularly in their C-terminal halves. Alpha-crystallin has chaperone-like properties including the ability to prevent the precipitation of denatured proteins and to increase cellular tolerance to stress. The modeled structure revealed N-terminal arm of the PgHSP16.97 represents an extensive, intrinsically unstructured domain rich in hydrophobic residues (53%) which will play key roles in protein– protein interactions with denatured proteins and thus critical to substrate interactions. Structural disorder allows the N-terminal arm to present a variable and flexible ensemble of clusters of hydrophobic residues that can interact with diverse geometries of hydrophobic patches on unfolding proteins [43]. This ability to present multiple binding site conformations makes PgHSP16.97 highly effective at interacting efficiently to protect a wide range of critical cellular proteins. Also N-terminal regions are important for stabilizing the oligomer through interlocking subunits by forming two disks intertwines to form pairs of knot-like structure, and the hydrophobic contacts in these knots are buried inside the oligomer [44]. The C-terminal extension is variable in length and its function in those cellular compartments enigmatic. The sequence information revealed the C-terminal extension was rich in glutamic acid residues (E-), which were critical for its chaperonic activity [45]. Also there was Ile-X-Ile residue at C-terminal extension (β 10 strand) which has a role in oligomerisation of heat shock proteins [46] by interacting with the hydrophobic pockets formed at β4 and β8 of ACD strands. There were two consensus regions within C-terminal separated by a region with hydrophilic residues forms the signature sequence for identification of cytosolic plant sHSP. The consensus region I (CR-I) consists of residues (P-X_14-_GVL) which is involved in multimerisation of PgHSP 17 subunits and Consensus region II (CR-II) consists of residues (P-X_15-_V-L) involved in solubility of protein complex. Arginine (R) conserved across the cereal sHSP gene at position 114 is responsible for stabilization of dimer by formation of intermolecular salt bride with Glutamic acid at position 100 [44]. A phylogenetic study was conducted using HSP *17.0*full length protein sequences belongs to C I sHSP family from seven different cereals, along with *Pg HSP 16.97* using Clustal W included in the MEGA 5.2 software (Fig.7). There is considerable diversity in *HSP 17.0* evolution in cereals particularly at N-terminus where the *Pg HSP 16.97* is showing close similarity to Pg HSP 17.0 (X94191.1) and is evolutionarily very close to Zea *HSP 16.9* protein and is more divergent to that of sorghum.

## Conclusions

Expression profiling becomes a powerful tool in identifying and classifying the genotypes carrying novel genes based on their expression upon specific condition or growth stages. In this study, an attempt was carried out to validate the powerfulness of expression pattern study as molecular screening techniques in identifying thermotolerant lines based on the expression of HSPs or HSFs genes at seedling stage of Pearl millet and bridging them together to fight against the unpredicted nature of abiotic stress. These results provide a comprehensive molecular biology background for research on thermo-tolerance among crop plants, particularly with respect to the structural and functional aspects of sHSPs. All of the genes undertaken in our research have significance for breeding Pearl millet with increased thermotolerance.

## Acknowledgments

This study was a part of M.Sc thesis and was supported by the DBT funded project on “Molecular cloning and functional characterization of *annexin* family genes from Pearl millet (*Pennisetum glaucum*) under abiotic stress”. The First author (MSS) wishes to thank ICAR-IARI, New Delhi for providing the facilities for the study and the JRF provided by ICAR, India.

## Author Contributions

Conceptualization and designing the experiments: CTS & SB.

Performed the experiments: MSS

Contributed reagents/materials/analysis tools: SPS, RK & SSL

Analyzed the data: SSL.

Wrote the original draft: MSS, CTS, SB

Review & editing: CB & SPS

Project administration: SB, CTS & KVP

